# DNA-free and genotype-independent CRISPR/Cas9 system in soybean

**DOI:** 10.1101/2024.04.02.587856

**Authors:** Chikako Kuwabara, Ryuji Miki, Nobuyuki Maruyama, Masanori Yasui, Haruyasu Hamada, Yozo Nagira, Ryozo Imai, Naoaki Taoka, Tetsuya Yamada

## Abstract

The CRISPR/Cas9 system has revolutionized the field of plant genetic engineering. Here we report a smart genome editing system of soybean by using iPB-RNP method without introducing foreign DNA and requiring traditional tissue culture processes such as embryogenesis and organogenesis. Shoot apical meristem (SAM) of embryonic axes was used the target tissue for genome editing, because the SAM in soybean mature seeds has stem cells and specific cell layer developing germ cells during reproductive growth stage. In the iPB-RNP method, the complex of ribonucleoprotein (RNP) and Cas9 protein was directly delivered into SAM stem cells via particle bombardment and genome-edited plants were generated from these SAMs. Soybean allergenic gene *Gly m Bd 30K*, which we previously generated genome-editing soybean by using *Agrobacterium*-mediated transformation and particle bombardment in our previous studies, was targeted in this study. Many E_0_ (the first generation of genome-edited) plants in this experiment harbored mutant alleles at the targeted locus. Editing frequency of inducing mutations transmissible to the E_1_ generation was approximately 0.4 to 4.6 % of all E_0_ plants utilized in various soybean varieties. Furthermore, Gly m Bd 30K protein in mature seeds was not detected by western blot analysis due to flame-sift mutations. Our results offer a practical approach for both plant regeneration- and DNA-free genome editing achieved by delivering RNP into the SAM of dicotyledonous plants.

## Introduction

The CRISPR/Cas9 system has revolutionized the field of plant genetic engineering^1-3^. To facilitate advanced and precise site-directed mutagenesis in plants, the expression module of the CRISPR/Cas9 system is frequently integrated into the host genome as a foreign DNA. This integration is typically achieved through methods such as particle bombardment or *Agrobacterium*-mediated transformation^4-5^. However, the process of genome editing often encounters challenges in specific target plant species and strains. These challenges stem primarily from limitations in the available genotypes for the crucial step of plant regeneration during the transformation process. Notably, methods like *Agrobacterium*-mediated floral dip in Arabidopsis^6^ or particle bombardment in wheat^7^ have been successful in directly producing genome-edited plants, circumventing the traditional process of plant regeneration. More recently, particle bombardment in wheat has facilitated DNA-free genome editing using the ribonucleoprotein (RNP) complex of the CRISPR/Cas9 system, known as the iPB-RNP method^8^. In this study, we optimized the iPB-RNP system for efficient genome editing in the shoot apical meristem (SAM) of soybean, representing a key dicotyledonous plant species. We observed that many E_0_ (the first generation of genome-edited) plants in this experiment harbored mutant alleles at the targeted genetic locus. Additionally, we found that the frequency of inducing mutations transmissible to the E_1_ generation was approximately 0.4 to 4.6 % of all E_0_ plants utilized in this study. Our research thus offers a practical approach for both plant regeneration- and DNA-free genome editing, achieved by delivering RNP into the SAM of dicotyledonous plants.

Two major transformation platforms, namely *Agrobacterium*-mediated and particle-bombardment methods, have become foundational in the realm of genome editing across various plant species^4-5^. However, these transformation processes, which are essential for establishing transgenic plants, are significantly influenced by the genotype of the host plant. This dependency is especially notable in the context of dedifferentiation, somatic embryogenesis, and organogenesis, which are vital steps for the establishment of transgenic plants and rely on the type of tissue culture and the organ used as an explant^9^. Interestingly, *in planta* transformation systems sometimes enable the generation of transgenic plants without necessitating traditional tissue culture processes such as embryogenesis and organogenesis. This transformative approach has also been applied in genome editing using the CRISPR/Cas9 system. The *Agrobacterium*-mediated floral dip transformation method has been successfully utilized to stably generate genome-edited Arabidopsis plants employing the CRISPR/Cas9 system^6^. In wheat, the SAM of immature seeds was used as the explant for plant transformation, employing the expression module of the CRISPR/Cas9 system^7^.

In practical applications of genome editing, the persistence of foreign DNA within the genome can limit the commercial release of edited crops and vegetables. If the expression module of the CRISPR/Cas9 system remains integrated in the host genome, there is a risk of inducing off-target mutations, which may result in unexpected traits in the transgenic plants^10-12^. Typically, foreign genes are eliminated from the host genome through genetic segregation in subsequent generations^13^. In some cases, transient expression of the CRISPR/Cas9 module via *Agrobacterium* has led to the production of mutated plants that do not contain the foreign DNA in the T_0_ generation, as observed in tobacco^14^. Viral vectors have also been effectively employed for DNA-free genome editing. For instance, the sonchus yellow net rhabdovirus-based viral delivery system of the CRISPR/Cas9 module has enabled efficient genome editing in *N. benthamiana*^15^. Moreover, these viral vectors can often be easily eliminated from mutant plants during the process of plant regeneration or subsequent seed setting^15^. It is noteworthy that these approaches generally require the step of plant regeneration to produce genome-edited plants^14-15^.

The direct delivery of RNP, consisting of gRNA and Cas9 protein, has been increasingly utilized for DNA-free genome editing in plants. The introduction of RNPs into the shoot apical meristem (SAM) on immature embryos via particle bombardment has led to the generation of DNA-free mutant plants in species such as maize^16^ and wheat^17^. These approaches, however, also necessitate the process of plant regeneration. More recently, the delivery of RNP into the SAM of mature embryos, using the *in planta* bombardment system (iPB), has allowed for the production of genome-edited wheat plants without the need for a regeneration process^8^. Nonetheless, the full potential of the iPB-RNP approach for dicotyledonous plants, such as soybeans, remains to be fully explored and optimized.

## Results

In our experimental setup, an embryonic axis was prepared from germinated soybean seeds of the Yukihomare variety (Fig. 1a). We removed the small primary unifoliolate leaves from the embryonic axis, thereby exposing the SAM at the top of the embryonic axis (Fig. 1b-d). The exposed SAM presented a single dome shape (Fig. 1e). To better understand the physiological and structural characteristics of the soybean SAM, we compared it with the wheat SAM, which has already been successfully genome-edited using the iPB-RNP method^8^. The soybean SAM, located between two unifoliolate leaf primordia (LP), had a significantly stiffer organization compared to the wheat SAM (Extended Data Fig. 1). In contrast, the wheat SAM at the germinating stage was covered by the second and third and fourth leaves and was damp and very tender compared to the soybean SAM (Extended Data Fig. 1). In comparison to wheat, the cell size in the soybean SAM (136.0 ± 18.0 μm^2^) was markedly smaller than wheat (288.7 ± 13.8 μm^2^) (n = 40-64, *P* < 0.001), and the soybean SAM consisted of a greater number of cells (Extended Data Fig. 1). Following the removal of the unifoliolate leaves, the soybean SAM formed a single LP within two days (Fig. 1f). Three days later, a total of three LPs had developed (Fig. 1g). Subsequently, trichomes began to cover the SAM (Fig. 1h, i), eventually leading to the SAM being completely enveloped by numerous trichomes (Fig. 1j). RNP bombardment was performed immediately after the SAM exposure. The delivery conditions of the RNP were determined based on the localization of GFP fluorescence using a Cas9-GFP fusion protein (Fig. 1k, l). These results indicate that the conditions for RNP delivery in soybean SAM are distinct from those in wheat (Supplemental Table 1).

**Table 1.** Genome editing of soybean by iPB-RNP method using 0.6 μm diameter gold particles.

| Exp. No. | No. of bombarded embryonic axis | No. of surviving explants | No. of mutants in bulked genomic DNA | No. of mutants in each genomic DNA | Frequency of genome editing (%) |
| --- | --- | --- | --- | --- | --- |
| 1 | 50 | 49 | 6 | 1 | 2.0 |
| 2 | 68 | 63 | 5 | 0 | 0 |
| 3 | 48 | 46 | 4 | 0 | 0 |
| 4 | 34 | 31 | 8 | 0 | 0 |
| 5 | 66 | 62 | 3 | 0 | 0 |
| 6 | 32 | 31 | 4 | 1 | 3.1 |
| 7 | 64 | 50 | 12 | 1 | 1.6 |
| 8 | 64 | 61 | 4 | 0 | 0 |
| 9 | 65 | 61 | 4 | 1 | 1.5 |
| 10 | 48 | 45 | 4 | 0 | 0 |
| 11 | 65 | 64 | 12 | 1 | 1.5 |
| Total | 604 | 563 | 66 | 5 | - |
| Ave. | 54.9 | 51.2 | 6.0 | 0.5 | 0.9 ± 0.3 |

Average of frequency of genome editing is expressed as the mean and standard error of 11 experiments.

**Fig. 1.**
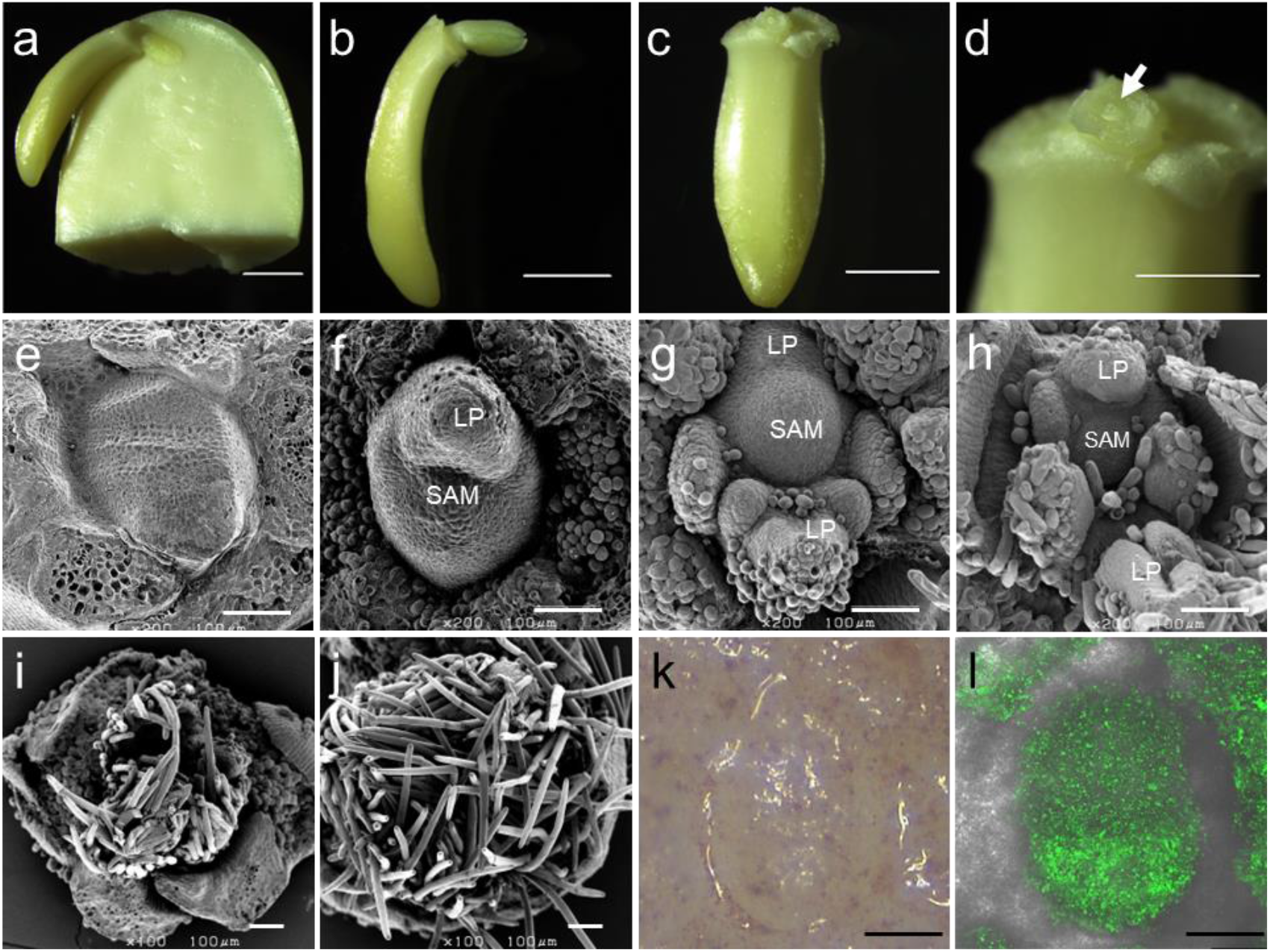
Visualization of soybean shoot apical meristem (SAM) and Cas9 protein delivery. **a-d**, Stages of Soybean Embryonic Axis: Cross-section of a mature seed’s embryonic axis post-imbibition (**a**). Isolated embryonic axis removed from cotyledons (**b**). Embryonic axis following the removal of primary unifoliolate leaves (**c**). SAM is located at the tip of the embryonic axis, as indicated by the arrow (**d**). Scale bars correspond to 2 mm (**a**-**c**) and 1 mm (**d**). **e-j**, Scanning Electron Microscopy of Shoot Apices: Images captured one to five days after imbibition. Day one post-imbibition of embryogenic axis (**e**), day two (**f**), day three (**g**), day four (**h** and **i**) with trichomes removed (**h**), and day five (**j**). LP denotes leaf primordia. Scale bars measure 100 μm. **k-l**, Confocal Laser Scanning Microscopy of Shoot Apices: Micrographs taken 24 hours post-iPB. Scale bar is 100 μm. Light micrograph showcasing the shoot apex (**k**). GFP fluorescence observed at the shoot apex (**l**).

The allergenic gene *Gly m Bd 30K*, which we previously modified using genome editing, was the focal point of this study as the targeted locus^18-19^. An illustrative overview of the RNP delivery and the subsequent schemes for analyzing mutagenesis is presented in Extended Data Fig. 2. Plantlets that emerged from the treated embryonic axes were designated as E_0_ plants. We analyzed mutations at the targeted locus in these E_0_ plants by employing cleaved amplified polymorphic sequences (CAPS) analysis on bulked genomic DNA extracted from several of their leaves (Fig. 2a, Extended Data Fig. 2). The DNA fragments resulting from this analysis were categorized into wild type and mutant types, based on the sizes of the fragments (Fig. 2a). Plants that originated from 66 of the explants, representing approximately 6% of the total explants used in this experiment, were identified as mutants (Table 1, Extended Data Fig. 2). Of these, five mutant E_0_ plants (Y19-102-1, Y19-134-10, Y19-253-13, Y19-250-15, and Y19-262-7) also displayed mutant alleles in the genomic DNA of each leaf (Table 1, Fig. 2b-c, Extended Data Fig. 2). The mutations identified in these alleles included both indels and nucleotide substitutions (Fig. 2c).

**Fig. 2.**
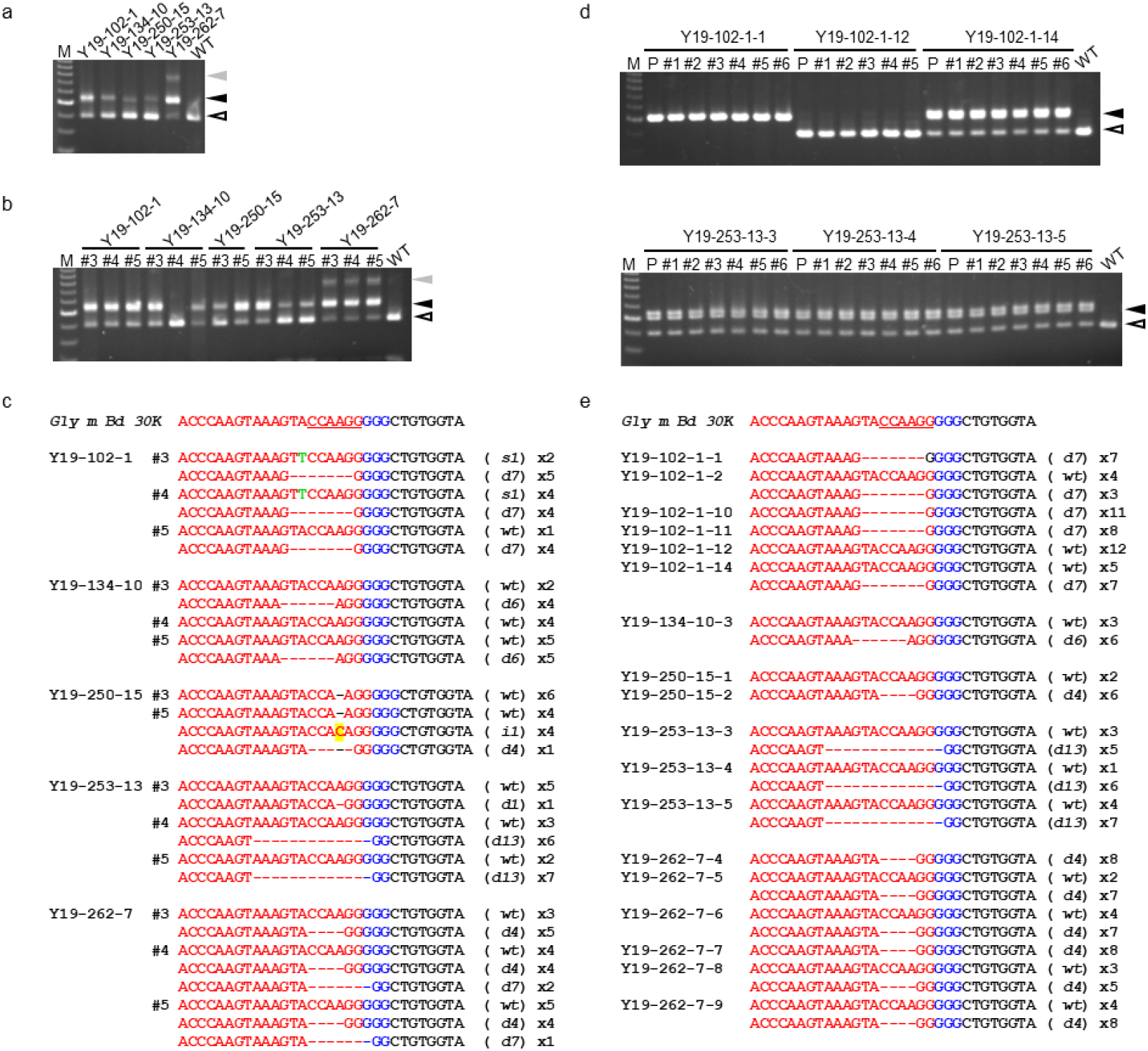
Mutation detection in E_0_ and E_1_ plants through CAPS and sequencing analyses. **a, b, d**, CAPS Analysis: CAPS analysis of bulked DNA from E_0_ leaves (**a**). Individual DNA analysis from E_0_ (**b**) and E_1_ (**d**) leaves. ‘#’ and P represent the leaf position and primary unifoliolate leaf, respectively. Branch numbers at the top of each lane indicate specific mutant plant numbers. Black arrowheads point to expected mutant-type fragments; white arrowheads indicate wild-type fragments; gray arrowheads show fragments of unexpected sizes. WT represents Yukihomare. M is the 100 bp ladder marker. **c, e**, Nucleotide Sequencing Analysis: Target locus and adjacent regions sequenced in E_0_ (**c**) and E_1_ (**e**) plants. *Gly m Bd 30K* corresponds to the Yukihomare sequence. Red and blue nucleotide sequences highlight the gRNA-targeted region and the proto-spacer adjacent motif (PAM) region, respectively. Mutation types are denoted in parentheses: e.g., *d1* (a single-nucleotide deletion); *i1* (a single-nucleotide insertion, highlighted in yellow); *s1* (a single-nucleotide restitution, green); *wt* (no mutation). ‘×’ indicates the number of sequenced clones. Sequences recognized by the BsaJ1 enzyme are underlined in *Gly m Bd 30K* sequences.

E_1_ seeds, harvested from these E_0_ mutants approximately 15 weeks after the bombardment process, were analyzed to confirm mutations at the targeted locus using the CAPS method (Extended Data Fig. 3). All but one line (Y19-250-15) exhibited a Mendelian segregation pattern of 1:2:1, corresponding to mutant homozygous, heterozygous, and wild type homozygous respectively (Supplemental Table 2). The spatial and temporal distribution of mutagenesis in both E_0_ plants and E_1_ seeds was established based on the CAPS analysis findings. Notably, E_1_ mutant seeds were exclusively obtained from nodes that had leaves where mutagenesis had been observed (Fig. 3). Altering the size of the gold particles or the frequency of bombardments in the genome editing process also resulted in the acquisition of mutant E_1_ seeds (Extended Data Fig. 4). Interestingly, the phenomenon of chimeric mutagenesis, which was observed in E_0_ plants, was not present in the E_1_ plants (Fig. 2d). Furthermore, the genotypes of mutant alleles in the E_1_ generation were consistent with those found in the E_0_ generation (Fig. 2e).

**Fig. 3.**
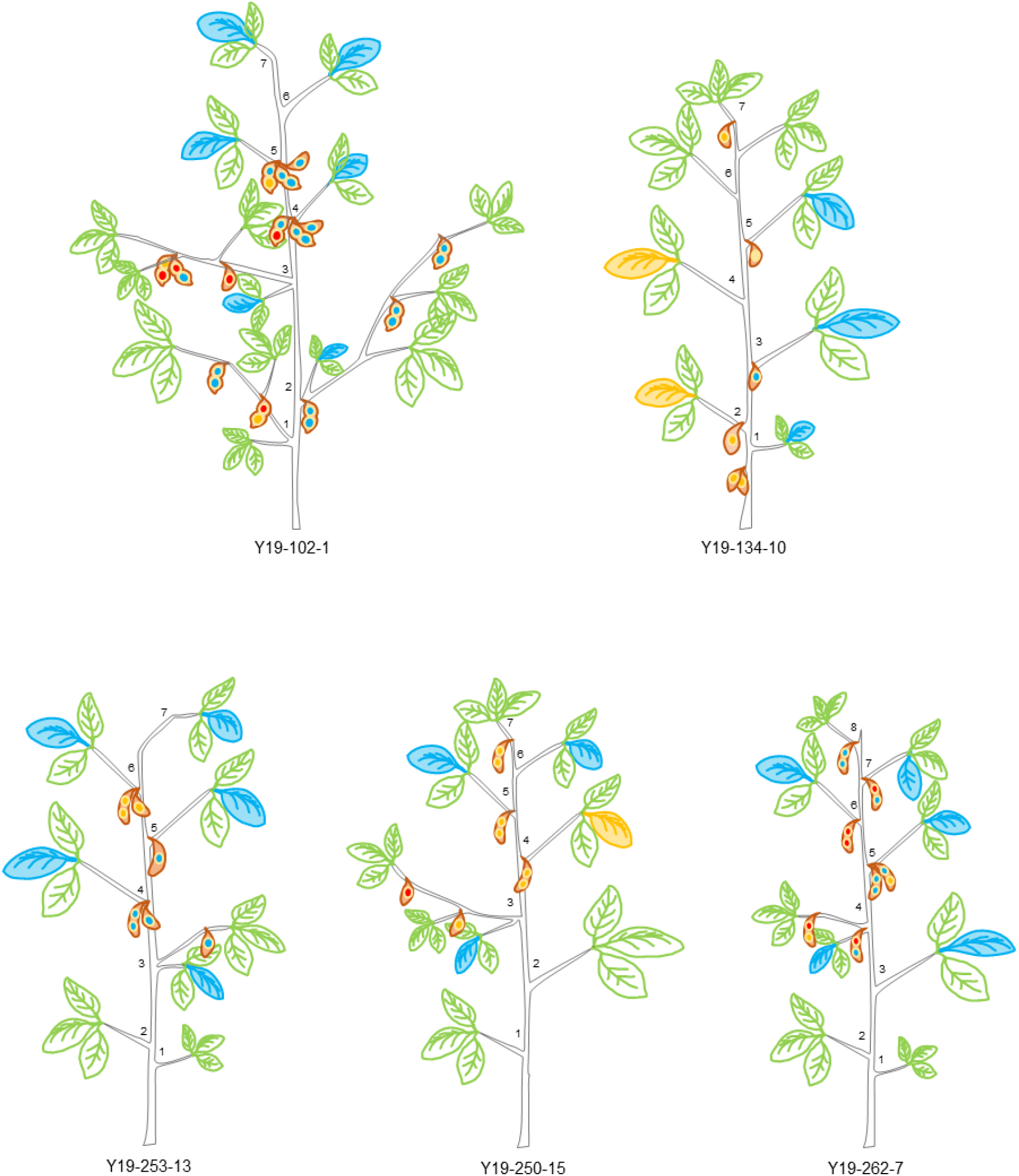
Spatial distribution of mutations in E_1_ plants and E_1_ seeds. The diagram indicates the positions of leaves on the stem, starting from the base of the E_0_ plant and excluding the primary unifoliolate leaf. The colors of leaves and seeds illustrate mutation presence or absence based on CAPS analysis results. Red signifies the presence of only mutant-type fragments; blue represents a mix of mutant- and wild-type fragments; yellow indicates only wild-type fragments.

The iPB-RNP system, having been optimized for soybean, was subsequently applied to two Japanese soybean varieties, Enrei and Fukuyutaka, as well as two US varieties, Williams82 and Jack. The mutation induction frequency transmitted to the E_1_ generation was approximately 0.4 to 4.6% of the total explants used in these cases (Supplemental Table 3). In the E_1_ seeds of these four varieties, frame-shift mutations were detected (Extended Data Fig. 5a). SDS-PAGE analysis revealed no significant differences in the protein composition of all mutant seeds, including those from the Yukihomare variety (Extended Data Fig. 5b). However, western blot analysis, conducted using a polyclonal antibody specific for Gly m Bd 30K, indicated a deficiency of this protein in mature seeds that had frame-shift mutations (Extended Data Fig. 5c).

## Discussion

The shoot apical meristem (SAM) is known to comprise three structurally distinct cell layers (L1 to L3)^20^. The L1 and L2 layers contribute to the formation of epidermal and subepidermal tissues, respectively^20^. Moreover, the L2 layer is intricately involved in the development of germ cells^21-22^. In wheat, for instance, the L2 layer is targeted in particle bombardment-mediated transformation and genome editing, as this enables the transmission of foreign genes to subsequent generations^7-8^. In our study, about 0.4 to 4.6% of the total embryonic axes resulted in mutant alleles at the targeted locus in the E_1_ generation (Table 1, Supplemental Table 3). This finding indicates that the RNPs were successfully delivered into the specific sector of the SAM responsible for giving rise to future germ cells. However, it is important to note that the region within the L2 layer where germ cells develop is limited. The SAM, both cytologically and functionally, is categorized into three zones: the central zone (CZ), the peripheral zone (PZ), and the rib zone (RZ), distinguished based on patterns of gene expression and function^23-25^. The CZ serves as a reservoir of stem cells, supplying cells to both the PZ and RZ, while also maintaining the integrity of the CZ zone itself^26^. Therefore, delivering RNPs precisely to the CZ within the L2 layer was essential. Although the depth of SAM layers reached by the RNP could be ascertained in our experiments using the GFP-Cas9 protein, the specific CZ region could not be clearly identified (Extended Data Fig. 2). Enhancing the efficiency of RNP delivery to the CZ region of the L2 layer could potentially increase the frequency of mutagenesis that can be transmitted to the next generation.

In this study, we occasionally detected two mutant alleles at the targeted locus in the E_1_ generation of plants grown from a single embryonic axis (Extended Data Fig. 4c and 5a). This observation suggests that if mutagenesis occurred independently at both alleles in a single stem cell destined to become a germ cell, then biallelic mutations should be observable in the E_1_ generation. Indeed, we identified E_1_ plants with biallelic mutations at the targeted locus (Extended Data Fig. 4c and 5a). This implies that two mutations were simultaneously induced in a stem cell shortly after the RNPs were delivered into it, and this cell was poised to generate germ cells. The stem cells that underwent mutagenesis at the targeted locus were also found to be involved in the differentiation of both leaf and reproductive tissues at each node of the E_0_ plant (Fig. 2). Therefore, confirming mutagenesis in the leaves of E_0_ plants served as an effective method for selecting mutants in the subsequent generation. However, it cannot be ruled out that the two mutant alleles may have originated from independently mutated cells that later differentiated into pollen and egg cells, respectively. Our findings demonstrate the iPB-RNP system’s potential for precise and rapid genetic improvement in various soybean genotypes.

## Materials and methods

### Preparation of soybean shoot apical meristem

Mature soybean seeds (*Glycine max* L. cv. Yukihomare, Enrei, Fukuyutaka, Williams 82, and Jack) were first sterilized using 1.5% sodium hypochlorite for 2 minutes and subsequently rinsed thoroughly, being washed ten times with sterile distilled water. These sterilized seeds were then soaked in sterile distilled water for 15-16 hours under dark conditions at a temperature of 25°C. For explant preparation, embryogenic axes were carefully excised from the cotyledons using a scalpel. Under a stereomicroscope, unifoliate leaves and primary stipules were delicately removed. A total of 16 explants were arranged in a circular pattern, each approximately 1 cm in diameter, on a bombardment medium. This medium comprised Murashige-Skoog (MS) medium (pH 5.7, Sigma-Aldrich), 30 g/L sucrose (FUJIFILM WAKO Pure Chemicals), 0.5 g/L MES (Dojindo Laboratories), 20 mg/L meropenem (Sumitomo Dainippon Pharma), and 4.5 g/L phytagel (Millipore Sigma), with the shoot apical meristem region oriented upwards.

### Preparation of gold particles and particle bombardment

The GFP-Cas9 fusion protein was sourced from the National Agriculture and Food Research Organization. Both the crispr RNA (crRNA) and *trans-*activating crRNA (tracrRNA) were procured from FASMAC Co., Ltd. The target DNA sequence for *Gly m Bd 30K* was designated as 5′-ACCCAAGUAAAGUACCAAGG-3′. To assemble a single guide RNA (sgRNA) and *Streptococcus pyogenes* Cas9 (SpCas9) complex, guide crRNA and tracrRNA, each at a concentration of 1 μg/μl, were mixed and incubated on ice for 10 minutes. Subsequently, SpCas9 protein (NEB), 1 x cut smart buffer (NEB), and RNase inhibitor (Thermo Fisher Scientific) were added and the mixture was incubated at room temperature for an additional 10 minutes. This sgRNA-SpCas9 complex solution was then combined with TransIT-LT1 reagent (Mirus Bio) and incubated at room temperature for 5 minutes. To this mixture, 0.6 μm gold particles (450 μg/shot, BioRad) were added and incubated on ice for 10 minutes. The suspension was then centrifuged at 1,200 x *g* for 1 second, and the resulting pellet was resuspended in nuclease-free water. The sgRNA/SpCas9-coated gold particle suspension was evenly distributed onto a hydrophilic membrane (Scotchtint, 3M) attached to a macrocarrier and allowed to dry naturally prior to bombardment.

Bombardment was executed using a Biolistic PDS-1000/He Particle Delivery System (Bio-Rad), following a slightly modified version of the instruction manual. The distance maintained between the target plantlets and the edge of the brass adjustable nest of the microcarrier launch assembly was approximately 3.5 cm. Each plate containing the explants was bombarded at a helium pressure of 1,800 psi. Post-bombardment, the explants were incubated for 24 hours at 25°C in darkness.

### Caring of explants

Twenty-four hours after bombardment, the explants were transferred to MS medium (pH 5.7) supplemented with 30 g/L sucrose, 0.5 g/L MES, 20 mg/L meropenem, 3 ml/L Plant Preservative Mixture (Plant Cell Technology), and 3 g/L phytagel. These explants were then maintained at a temperature of 25°C under an 18-hour light photoperiod at an intensity of 16 μmol m^-2^ s^-1^. Approximately two weeks later, explants that had successfully elongated shoots and roots were individually relocated to seeding trays filled with potting soil (Sumitomo Forestry Landscaping). To prevent desiccation of the young leaves, each tray was covered with a plastic sheet. The plantlets were maintained at 25°C under a 16-hour light photoperiod at 50 μmol m^-2^ s^-1^ until they reached a height of about 8 cm. The E_0_ plants were subsequently transferred to plastic pots filled with a mixture of potting soil for sowing and culture soil (Katakura & Co-op Agri Corp) in a 1:3 ratio. They were then kept in a closed greenhouse under a 12-hour photoperiod at 25°C until the E_1_ seeds were collected.

### Detection of genome edited plants

Genome-edited E_0_ and E_1_ plants were identified using cleaved amplified polymorphic sequences (CAPS) analysis. Genomic DNA was isolated from several leaves of each bombarded plant, ensuring that these leaves were sourced from varying positions. To amplify the targeted region, *Gly m Bd 30K* specific primers, 5′-GCAAGCTCCCAAGGATGTG-3′ and 5′-ACGCCCAACCGCTTCCTAT-3′, were utilized for PCR. The PCR reaction, facilitated by ExTaq polymerase (TaKaRa), was conducted in a 10 μl volume. This process commenced with an initial denaturing step at 95°C for 5 minutes, followed by 30 cycles of amplification: denaturing at 95°C for 30 seconds, annealing at 64°C for 30 seconds, and synthesis at 72°C for 30 seconds, culminating in an extension phase at 72°C for 7 minutes. Post-amplification, the DNA was subjected to digestion using the restriction enzyme BsaJ1 (NEB) at 60°C for 90 minutes, then at 80°C for 20 minutes. The digested DNA fragments were subsequently analyzed through 2% agarose gel electrophoresis and visualized with ethidium bromide staining under UV light.

### Sequencing analysis

To further investigate mutations at the target locus, PCR products from the CAPS analysis were cloned into the pGEM T-easy vector (Promega) and purified using ExoSAP-IT reagent (Affymetrix). Sequencing of these PCR products was performed on an ABI3130 sequencer (Applied Biosystems). This sequence analysis was carried out by the Instrumental Analysis Division at the Graduate School of Agriculture, Hokkaido University, and occasionally by FASMAC Co., Ltd. through their contract analysis service.

### Protein analyses of E_2_ mature seeds

The process for extracting crude seed proteins and conducting protein analysis was performed as described by Adachi et al. (2021).

### Microscopy

For scanning electron microscopy (SEM), shoot apices immersed in water for 1 to 7 days were fixed overnight in formalin-acetic acid-alcohol (FAA). These samples were then dehydrated through an ethanol series ranging from 50% to 100%. The dehydration process was followed by drying with a critical point dryer (EM CPD300, Leica Microsystems) and subsequent gold-palladium evaporation using a magnetron sputterer (MSP-20-MT, Vacuum Device) as per the manufacturer’s instructions. The prepared samples were observed using a scanning electron microscope (JSM-6301F, JEOL). The presence of Cas9 in the shoot apical meristem was investigated by observing the fluorescence of the Cas9-GFP protein (Venus-Cas9) using a confocal laser scanning microscope (TCS SP5, Leica Microsystems). The area of each cell within the SAM was calculated using SEM images of the SAM with Image J software (https://imagej.nih.gov/ij/download.html).

### Statistical analysis

For statistical purposes, 10 to 21 cells from four independent SAMs were selected randomly. Data analysis was conducted using the *t*-test with R version 4.2.1 (http://www.r-project.org/), considering *P* < 0.05 as statistically significant. To assess the fit between observed genetic segregation and the 1:2:1 segregation ratio hypothesis, a chi-square test was performed using Microsoft Excel, also setting a significance level of *P* < 0.05.

## Acknowledgements

We thank Motomi Suzuki, Sayuri Noguchi, Yasuko Kitsui, and Takako Yamamoto for general technical assistance. This work was supported by the Cross-Ministerial Strategic Innovation Promotion Program (SIP; Council for Science, Technology and Innovation, Cabinet Office, Japan) and the administration of individual commissioned project study (Development of new varieties and breeding materials in crops by genome editing, the Ministry of Agriculture, Forestry and Fisheries, Japan).

## Author contributions

CK, RM, YN, and TY designed the study. CK, RM, NM, MY, HH, YN, and TY contributed to the data collection and analysis. RI, NT, and TY contribute the data discussion and conceived the study and wrote the paper.

