## Supplemental file for "DNA-free and genotype-independent CRISPR/Cas9 system in soybean"

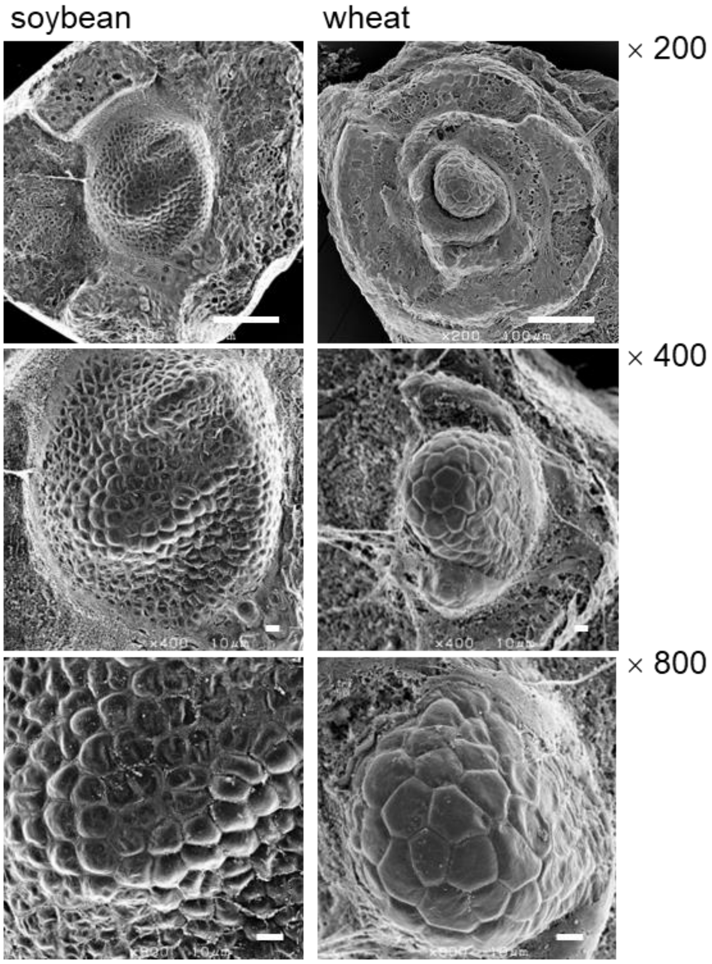


**Extended Data Fig. 1 Comparison of SAM characteristics in soybean and wheat. Scanning electron micrographs of vegetative shoot apices of soybean (left panels) and wheat (right panels) at 1 day after imbibition. Scale bars indicate 100 µm (× 200), 10 µm (× 400) and 10 µm (× 800).**

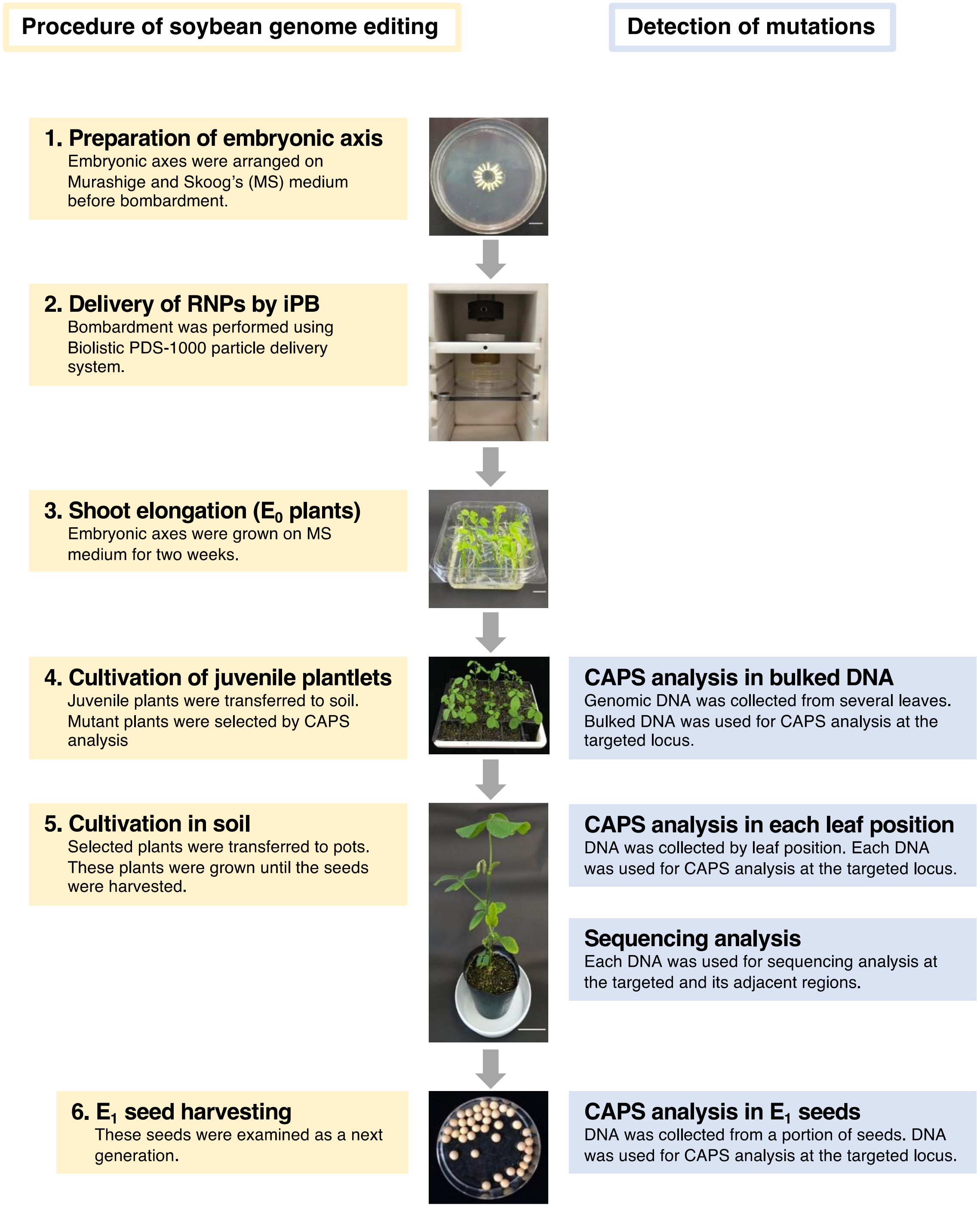


**Extended Data Fig. 2 Procedure by soybean genome editing by iPB-RNP method.**

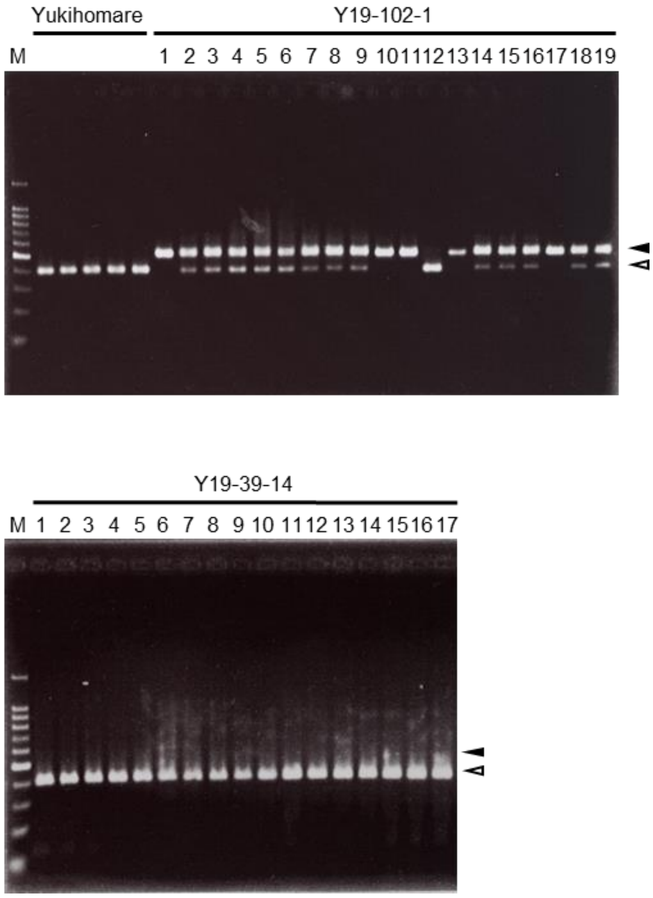


**Extended Data Fig. 3 Detection of mutations in $E_1$ seeds by CAPS analysis. Each lane in the analysis is labeled with a specific number, denoting the $E_1$ seed number. Additionally, at the top of each lane, numbers accompanied by branch symbols represent the corresponding mutant plant numbers. Within the analysis, different types of DNA fragments are indicated by arrowheads in various colors: black arrowheads point to the expected mutant-type fragments, while white arrowheads identify wild-type fragments. In this context, 'WT' refers to the wild-type control sample. The marker 'M' signifies the 100 base pair (bp) ladder marker, which serves as a reference for estimating the sizes of DNA fragments in the gel.**

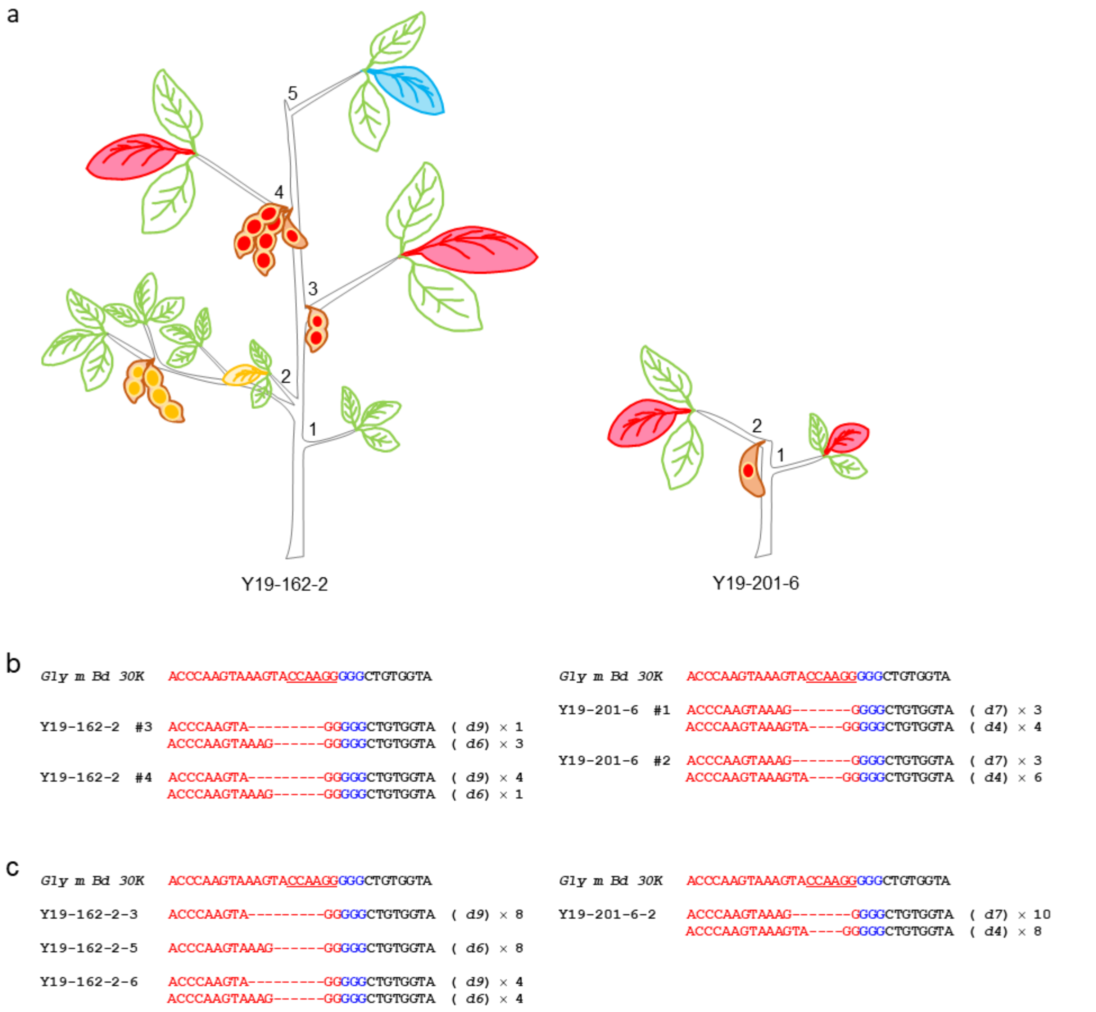


**Extended Data Fig. 4 Detection of mutations in $E_0$ plants and $E_1$ seeds. Spatial distribution of mutations in $E_0$ plants and $E_1$ seeds (a). The mutants depicted were generated under different iPB conditions compared to those in Fig. 3. Specifically, mutant Y19-162-2 was produced using 1.0 µm diameter gold particles, and mutant Y19-201-6 underwent bombardment at 1,800 psi twice. The numbers marked along the stem represent the leaf positions, counting upwards from the base of the $E_0$ plant. The color coding of leaves and seeds indicates the presence or absence of mutations, as determined by CAPS analysis. Red represents leaves or seeds with only mutant-type fragments, blue denotes the presence of both mutant- and wild-type fragments, and yellow signifies only wild-type fragments. Nucleotide sequencing analysis of the targeted locus and its flanking region. Sequences in $E_0$ (b) and $E_1$ (c) plants. The symbol '#' is used to indicate the leaf position. The nucleotide sequence of *Gly m Bd 30K* corresponds to the Yukihomare sequence. Within these sequences, the regions targeted by the guide RNA (gRNA) are highlighted in red, and the proto-spacer adjacent motif (PAM) regions are marked in blue. Mutations are identified with letters and numbers in parentheses, for example, '*d4*' indicates a deletion of four nucleotides. The notation '×' followed by a number refers to the count of sequenced clones. Nucleotides that are recognized by the BsaJ1 enzyme within the *Gly m Bd 30K* sequences are underlined for emphasis.**

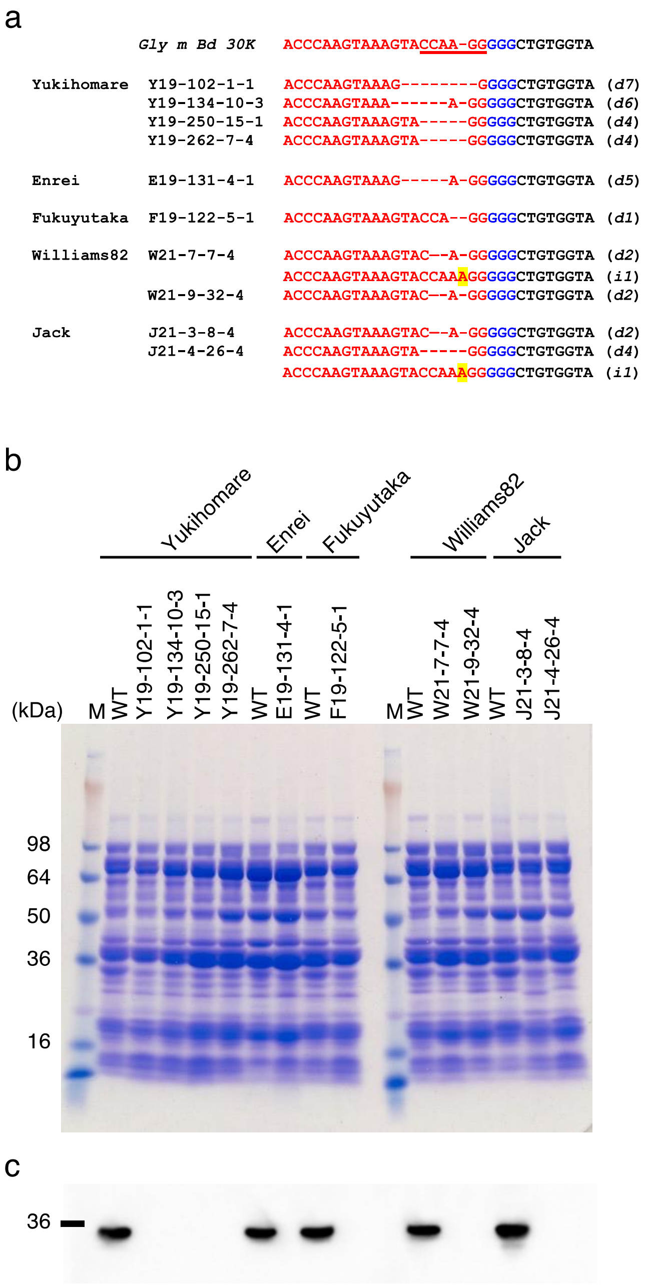


**Extended Data Fig. 5 Analysis of Gly m Bd 30K protein in mutant seeds. Nucleotide sequencing analysis of the targeted locus in $E_2$ and $E_1$ Seeds: This analysis focuses on $E_2$ seeds (from Yukihomare, Enrei, and Fukuyutaka mutants) and $E_1$ seeds (from Williams82 and Jack mutants). (a) The reference sequence, *Gly m Bd 30K*, is used to compare the gRNA-targeted region (red) and the proto-spacer adjacent motif (PAM) region (blue). Mutation types, such as '*d4*' for a four-nucleotide deletion and '*i1*' for a single-nucleotide insertion (highlighted in yellow), are identified. Nucleotides recognized by the BsaJ1 enzyme in the *Gly m Bd 30K* sequences are underlined. SDS-PAGE (b) and immunoblot analysis (c) were conducted on crude proteins extracted from wild-type and mutant**

**seeds ($E_2$ mutants from Yukihomare, Enrei, and Fukuyutaka; $E_1$ mutants from Williams82 and Jack). These analyses help compare protein profiles and detect the presence of the Gly m Bd 30K protein in both wild-type and mutant seeds.**

**Supplementary Table 1.** Comparison of genome editing experiment with soybean and wheat by *in planta* particle bombardment.

| Plant | Gold particle size (diameter, µm) | Gold particle content (µg/bombardment) | Pressure of bombardment (psi) | Times of Bombardment | Distance* (cm) |
|---|---|---|---|---|---|
| Soybean | 0.6 | 450 | 1,800 | 1 | 3.5 |
| Wheat | 0.6 | 270 | 1,350 | 4 | 6.0 |

*The distance indicates the length from the stopping screen to SAM of explants.

**Supplementary Table 2.** Number of $E_1$ seeds from each $E_0$ plant and genotyping of the *Gly m Bd 30K* gene by CAPS analysis.

| $E_0$ plant number | Total number of $E_1$ seeds | CAPS analysis pattern of $E_1$ seeds | | | $c^2$ | *P*-value |
|---|---|---|---|---|---|---|
| | | Mutation | | Wild type | | |
| | | Hetero | Homo | Homo | | |
| Y19-102-1 | 25 | 17 | 5 | 3 | 3.56 | 0.17 |
| Y19-134-10 | 5 | 1 | 0 | 4 | 8.2 | 0.02 |
| Y19-250-15 | 8 | 0 | 1 | 7 | 17 | 0.0002 |
| Y19-253-13 | 8 | 5 | 0 | 3 | 2.75 | 0.25 |
| Y19-262-7 | 14 | 7 | 5 | 2 | 1.29 | 0.53 |

Hetero of mutation indicates that both mutant and wild types are present.

**Supplementary Table 3.** Frequency of genome editing in Enrei, Fukuyutaka, Williams 82, and Jack by iPB-RNP method.

| Cultivar | Bombarded plants | Survival Plants | CAPS positive plants in $E_0$ progeny | CAPS positive plants in $E_1$ progeny | Frequency of genome editing (%) |
|---|---|---|---|---|---|
| Enrei | 160 | 156 | 6 | 1 | 0.6 |
| Fukuyutaka | 225 | 200 | 14 | 1 | 0.4 |
| Williams 82 | 191 | 162 | 19 | 9 | 4.6 |
| Jack | 128 | 127 | 6 | 2 | 1.5 |
